# Mechanism of nascent chain removal by the Ribosome-associated Quality Control complex

**DOI:** 10.1101/2025.01.03.631235

**Authors:** Wenyan Li, Talia Cahoon, Peter S. Shen

## Abstract

Errors during translation can cause ribosome stalling, leaving incomplete nascent chains attached to large ribosomal subunits^1–5^. Cells rely on the Ribosome-associated Quality Control (RQC) complex to recognize, process, and remove these partially synthesized proteins to maintain proteostasis. Despite its central role, the mechanisms by which the RQC complex orchestrates nascent chain processing and extraction have remained unclear. Here, we present a cryo-EM structure of the RQC complex from budding yeast, revealing how its core eukaryotic components function in nascent chain removal. We show that the Cdc48 protein extracting enzyme, along with its Ufd1-Npl4 adaptor, is recruited by the Ltn1 E3 ubiquitin ligase to extract ubiquitylated stalled peptides from the 60S ribosome. Additionally, we find that the Rqc1 component bridges the 60S ribosome with ubiquitin and Ltn1, facilitating the specific formation of K48-linked polyubiquitin chains on stalled peptides. These findings provide a structural and mechanistic framework for understanding how components of the RQC complex work together to remove faulty translation products from the ribosome, advancing our understanding of cellular protein quality control.

## Main

All cells depend on accurate protein synthesis to maintain proteostasis. From bacteria to humans, errors during translation can cause ribosome stalling, leading to the production of incomplete, potentially proteotoxic nascent chains and dissociation of the large and small ribosomal subunits^1–5^. The large subunit retains the aberrant tRNA-conjugated nascent chain, which is recognized, modified, and removed by the Ribosome-associated Quality Control (RQC) complex for its subsequent degradation. Mutations in RQC components are linked to neurological diseases, underscoring the importance of this pathway in maintaining protein homeostasis^6–9^.

In eukaryotes, the RQC complex consists of several conserved components, including Ltn1 (RING domain E3 ubiquitin ligase), Rqc2 (CAT tail synthase), Rqc1 (function unknown), and the Cdc48-Ufd1-Npl4 complex (protein segregase). The assembly of the RQC complex involves several key steps following the dissociation of 80S ribosomes. Initially, Rqc2 detects and binds to the obstructed large 60S ribosomal subunits containing peptidyl-tRNA. Rqc2 then recruits tRNAs charged with alanine and threonine to elongate the C-terminus of the nascent chain with CAT tails^10–17^. Rqc2 also facilitates the recruitment of Ltn1, which catalyzes the addition of K48-linked polyubiquitin chains onto the nascent polypeptide via its C-terminal RING domain^18,19^. Rqc1 plays a role in the formation of K48-linked ubiquitin chains by Ltn1 and is necessary to recruit Cdc48 to the stalled ribosome through an unknown mechanism^20–22^. The Cdc48-Ufd1-Npl4 complex is subsequently recruited to the stalled ribosome and presumably extracts the ubiquitylated nascent chain for proteasomal degradation^22–24^.

Despite extensive structural and functional studies of the RQC complex, many key steps underlying its mechanisms remain unresolved, including the role of the poorly understood but eponymous subunit, Rqc1, and the mechanism of stalled nascent chain extraction by Cdc48-Ufd1-Npl4. In this study, we determined cryo-EM structures of the fully-assembled RQC complex, providing a complete understanding how stalled nascent chains are K48 polyubiquitylated, extracted by Cdc48, and prepared for proteasomal degradation.

## Architecture of the eukaryotic RQC complex

RQC particles were isolated from budding yeast lysates through a rapid one-step affinity purification, as performed previously^10,22,25^. Cells expressed a C-terminal 3xFLAG tag at the endogenous locus of Rqc1, and the complex was purified by co-immunoprecipitation with anti-FLAG affinity resin (Extended Data Fig. 1). To enrich for Cdc48-bound particles, purifications were supplemented with the non-hydrolyzable ATP analog ADP-BeF_x_, which traps Cdc48 in its active, substrate-engaged conformation^26^.

Cryo-EM image processing revealed a subset of particles containing known and novel features associated with 60S particles (Fig. 1 and Extended Data Fig. 1 and 2). The reconstruction is consistent with previous studies that show Rqc2 binding to P- and A-site tRNAs at the exposed 60S-40S interface^10,11^. Ltn1 contacts Rqc2 at the sarcin-ricin loop and extends around the 60S ribosome to position its catalytic RING domain in the vicinity of the stalled nascent chain. Nascent chain density is observed within the exit tunnel and remains connected to the Rqc2-bound P-site tRNA. In addition to these previously established features of the complex, we resolved uncharacterized densities outside of the nascent chain exit tunnel through local refinement. The resulting maps enabled us to place models of Cdc48-Ufd1-Npl4 and Rqc1 in relation to the nascent chain and revealed the complete architecture of the RQC complex.

**Fig. 1.**
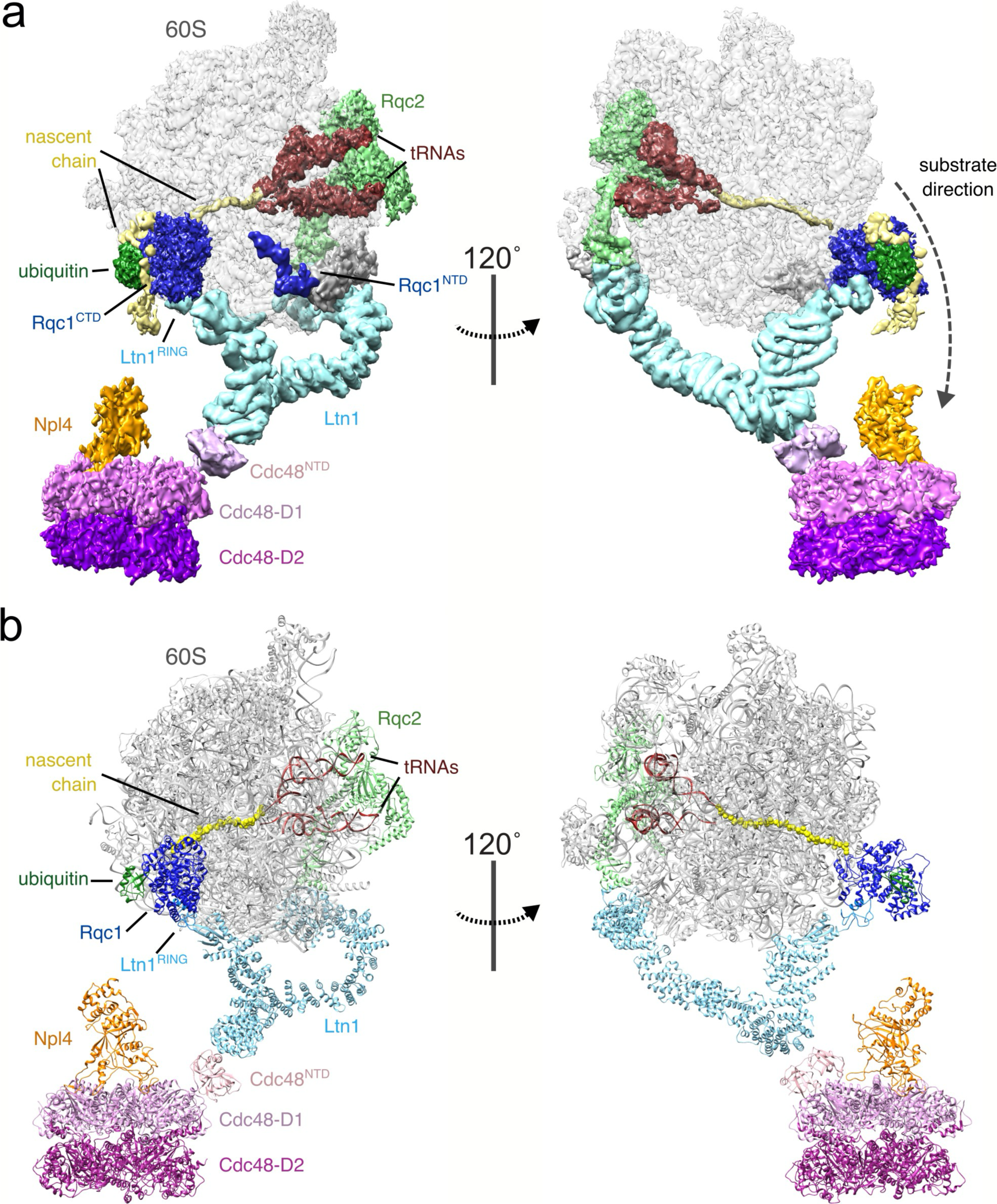
Architecture of the RQC complex. **a**, Composite map of the RQC complex segmented according to their individual components. **b,** Models of RQC components fitted into reconstruction density.

## Cdc48 interacts with Ltn1 on stalled ribosomes

The Cdc48 AAA+ ATPase is an abundant and essential enzyme, known for its role as a segregase across multiple cellular contexts. Cdc48 subunits comprise an N-terminal domain (Cdc48^NTD^), followed by tandem AAA+ motor domains, D1 and D2, which drive substrate unfolding. Structures of Cdc48 and its human ortholog p97/VCP show that substrates are unfolded by threading them through homo-hexameric assemblies of stacked D1 and D2 rings^26–29^. Cdc48 recruitment to specific subcellular localizations and substrates is regulated by its interactions with adaptor proteins, most notably the Ufd1-Npl4 heterodimer. Ufd1-Npl4 binds to K48-linked polyubiquitin chains and feeds substrates into the Cdc48 hexameric central pore to facilitate unfolding^30,31^. Despite their flexible association, our composite reconstruction places the entire Cdc48-Ufd1-Npl4 complex outside of the exit tunnel (Fig. 1 and 2). Focused particle sub-extraction and 2D classification over the density revealed characteristic features of the complex, including the stacked D1 and D2 ATPase rings and a tower-like density belonging to Ufd1-Npl4, which protrudes from the center of the particle^27,32^ (Fig. 2a and Extended Data Fig. 2). The peripheral Cdc48^NTD^ domains are positioned in the elevated, ‘up’ positions, which are associated with its active state^33^. Ufd1-Npl4 is positioned near the peptide exit tunnel and appears poised to transfer the ubiquitylated nascent chain from the 60S ribosome to Cdc48 (Fig. 1).

**Fig. 2.**
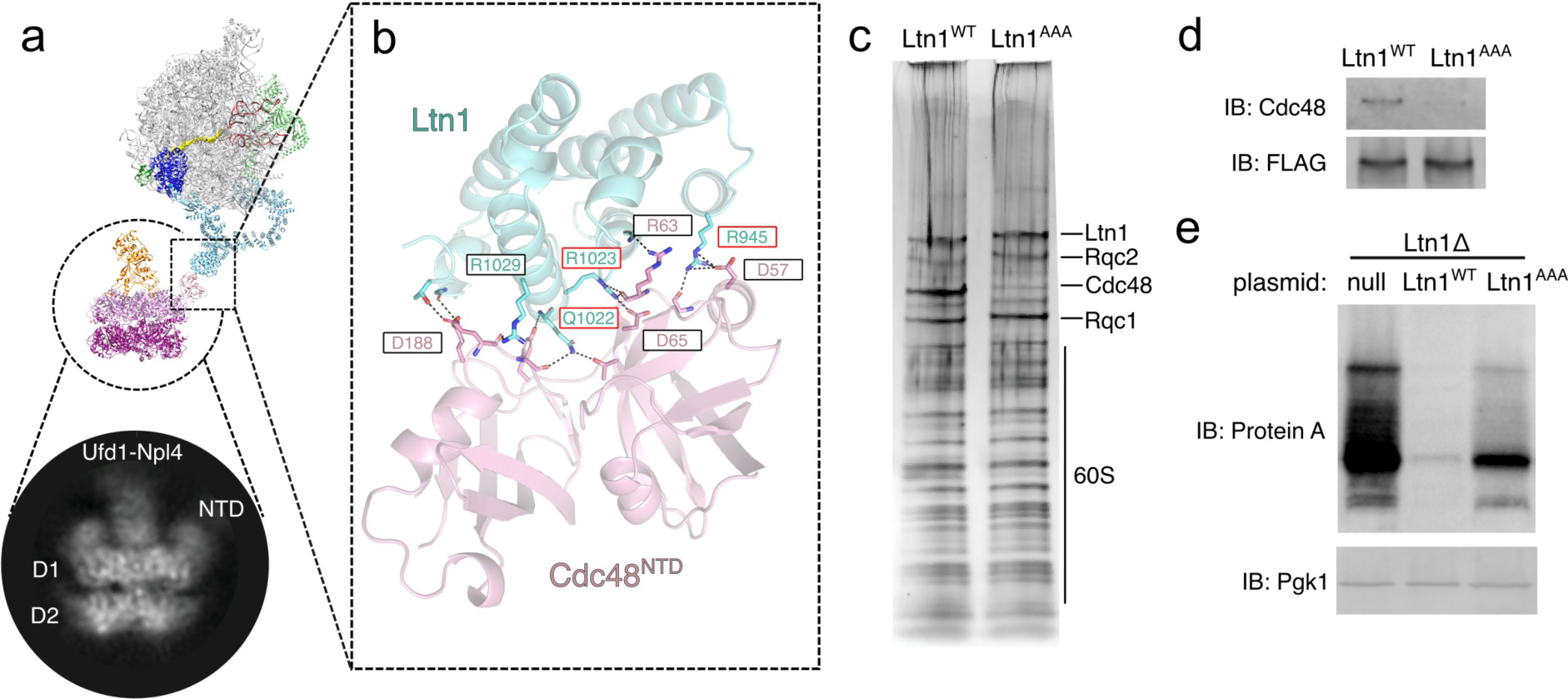
Cdc48-Ufd1-Npl4 is recruited to stalled ribosomes by Ltn1’s elbow domain. **a,** Positioning of Cdc48-Ufd1-Npl4 relative to the 60S ribosome. Focused view on bottom shows a 2D class average of sub-extracted Cdc48 complexes from RQC particles. **b,** Close up view of the Ltn1-Cdc48 interface at the Ltn1 elbow. Key interaction residues are labeled. Residues mutated to generate the Ltn1^AAA^ mutant are indicated with a red outline. **c,** Silver stained SDS-PAGE of RQC co-IPs expressing WT or Ltn1^AAA^. **d,** Anti-Cdc48 immunoblot from RQC co-IPs. Anti-FLAG (Rqc1) blot shown as loading control. **e,** Immunoblot of whole cell lysates expressing nonstop substrate (Protein A-nonstop) in Ltn1Δ cells and re-expressed with either wild-type Ltn1 or Ltn1^AAA^ mutant. Anti-Pgk1 blot shown as loading control.

The positioning of Cdc48-Ufd1-Npl4 relative to the 60S is facilitated through direct interactions between Cdc48^NTD^ and Ltn1. The interaction interface is located at the elbow of Ltn1, where its extended structure makes a sharp bend to position its catalytic RING domain in the vicinity of the emerging nascent chain. Focused 3D refinement over Ltn1 revealed sufficiently resolved density to place an AlphaFold3 model of the Ltn1-Cdc48^NTD^ interface as a rigid body (Fig. 2b and Extended Data Fig. 3a, b). Residues within four α-helices of Ltn1 form a network of interactions with the N-terminal double φ-barrel and a C-terminal β-barrel sub-domains of Cdc48^NTD^. Based on the predicted interactions, we performed mutagenesis experiments to test the relevance of the model’s contacting residues. Ltn1 residues R945, Q1022, and R1023 were mutated to alanines on an expression plasmid (R945A Q1022A & R1023A, Ltn1^AAA^) and transformed into Ltn1Δ cells. RQC co-IPs showed a significant reduction of Cdc48 recovery from cells expressing Ltn1^AAA^ compared to wild-type control, while other components of the RQC complex were recovered at similar levels (Fig. 2c, d). To test the functional relevance of the Cdc48-Ltn1 interaction, cells were expressed with a non-stop mRNA substrate and probed for substrate expression^34^. As expected, Ltn1Δ cells displayed an accumulation of non-stop substrate, and the degradation phenotype was rescued by re-expression of wild-type Ltn1 on a plasmid (Fig. 2e). In contrast, expression of Ltn1^AAA^ in Ltn1Δ cells led to an accumulation of stalled substrate at a milder extent compared to Ltn1Δ. Together, these experiments demonstrate that the Ltn1 elbow evolved to recruit Cdc48 to the dissociated 60S particles, and the loss of Cdc48 recruitment partially disrupts substrate degradation.

The direct interaction between Cdc48^NTD^ and Ltn1 establishes that Ltn1 is a Cdc48 adaptor. These adaptors typically contain conserved domains or motifs that define their interaction sites with Cdc48. The Ltn1 helical array interacts with Cdc48^NTD^ in a way that is reminiscent of the VIM and VBM motifs, where an α-helix sits in the groove that separates the φ-barrel and β-barrel subdomains^35^ (Extended Data Fig. 3c). However, unlike VIM and VBM motifs, which utilize a single α-helix for this interaction, Ltn1 does not share sequence similarity with these motifs and instead uses multiple helices to form a more extensive interface. Several E3 ligases are known to interact directly with the human ortholog of Cdc48 (p97/VCP), including HOIP^36^, gp78^37^, and HRD1^38^. These interactions likely drive Cdc48 localization to target 60S particles and couple substrate ubiquitylation directly to their unfolding.

## Rqc1 interacts with Ltn1 and ubiquitin

Rqc1 is the least characterized component of the RQC complex despite it being the source of co-IP experiments used for structure determination in previous studies^10,11^. Previous RQC structures lack Rqc1, presumably due to its dissociation from the particle during cryo-EM specimen preparation. Our cryo-EM specimens prepared using graphene grids likely shielded particles from the hydrophobic air-water interface and preserved Rqc1 on 60S particles. Cryo-EM reconstructions revealed uncharacterized densities ascribed to Rqc1 positioned near the opening of the nascent chain exit tunnel. The density is connected to ES27 of the 25S rRNA (nucleotides 1956-2092), Rpl38, and the Ltn1 RING domain (residues 1507-1562, Ltn1^RING^) (Fig. 3 and Extended Data Fig. 4 a-e). The quality of the density associated with Ltn1^RING^ was sufficiently resolved to fit the AlphaFold predicted structure of the Rqc1 C-terminal domain (CTD) as a rigid body (residues 176-682, Rqc1^CTD^) (Fig. 3a). By contrast, the other Rqc1 densities displayed poorer local resolution likely due to disorder. In support of this, the first 175 residues of Rqc1 (Rqc1^NTD^) are predicted to be mostly disordered, although we identify structured density corresponding to an internal helix (residues 122-135) that interacts with the 60S protein Rpl38 (Extended Data Fig. 4d, e). Despite the predicted disorder, the Rqc1^NTD^ is enriched with polar and basic residues that would be favorable to interact with rRNA and is consistent with a cloud of unstructured density associated with ES27 following the internal helix (Extended Data Fig. 4e).

**Fig. 3.**
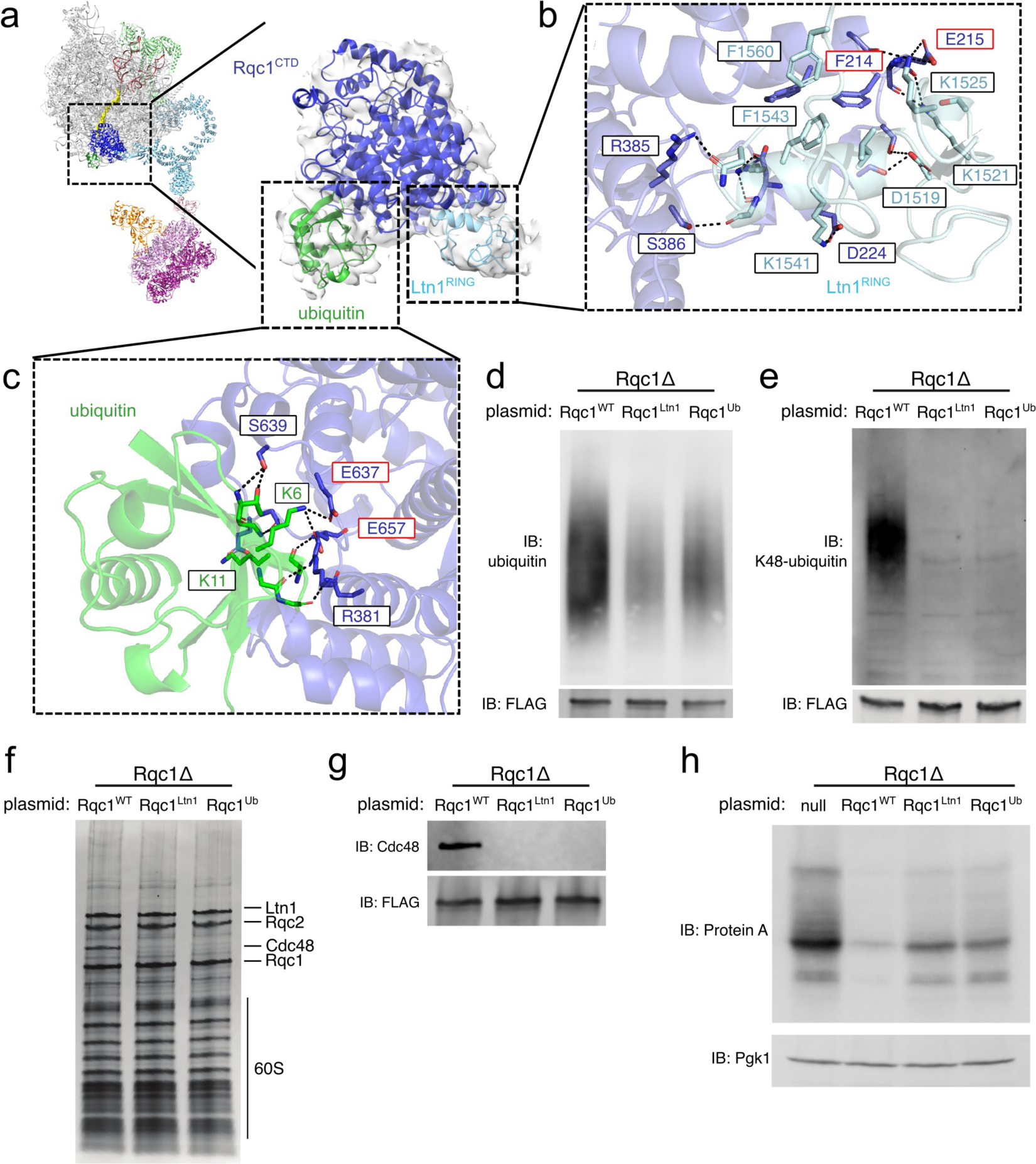
Rqc1 interacts with Ltn1 and ubiquitin. **a,** Close up view of Rqc1^CTD^ in complex with ubiquitin and Ltn1^RING^. **b, c,** Focused views of Rqc1 interactions with Ltn1^RING^ and ubiquitin, respectively. Key interaction residues are labeled. Residues mutated to generated Rqc1^Ltn1^ or Rqc1^Ub^ are indicated with a red outline. **d,** Immunoblot of RQC co-IPs from wildtype or Rqc1 mutant (Rqc1^Ltn1^ or Rqc1^Ub^) backgrounds probed against total ubiquitin. Anti-FLAG (Rqc1) blot shown as loading control. **e,** Same as in d, but probed with anti-K48-linked poly-ubiquitin antibody. **f,** Silver stained SDS-PAGE of RQC co-IPs reveal a loss of Cdc48 recovery by Rqc1^Ltn1^ and Rqc1^Ub^ mutants. **g,** Anti-Cdc48 immunoblot from Rqc1 co-IPs. Anti-FLAG (Rqc1) blot shown as loading control. **h,** Immunoblot of whole cell lysate against nonstop substrate (anti-Protein A) in Rqc1Δ cells expressing Rqc1^Ltn1^ or Rqc1^Ub^ mutants compared to wild-type control. Anti-Pgk1 blot shown as loading control.

The structure of Rqc1 suggests a bipartite mode of interaction such that the internal helix separates the Rqc1^NTD^ and Rqc1^CTD^ to facilitate interactions with the 60S ribosome and Ltn1^RING^, respectively. To test this model, we expressed a construct of Rqc1 that only contains its CTD, i.e. by deletion of residues 1-175. Expression and co-IP of this construct in an Rqc1 deletion strain (Rqc1Δ) failed to recover the 60S or other RQC components, thereby indicating that the Rqc1^CTD^ alone is insufficient for ribosome binding and RQC function (Extended Data Fig. 4f). Interestingly, ES27 and Rpl38 are also accessible on intact 80S ribosomes and would suggest that Rqc1 is capable of binding 80S particles as suggested in a previous study^20^. However, the specificity for 60S particles is augmented by the interaction between Rqc1 and Ltn1^RING^ on dissociated ribosomes.

Additional reconstruction densities were observed connected with Rqc1^CTD^ that were fitted well by models of its predicted interactions with Ltn1^RING^ and a ubiquitin molecule (Fig. 3a-c and Extended Data Fig. 4a). Modeling suggested that residues F214 and D215 of Rqc1 are crucial for its interaction with Ltn1^RING^ (Fig. 3b). We therefore substituted F214 and D215 with alanine to test how disruption of this interface would affect RQC function (F214A & D215A, Rqc1^Ltn1^). Immunoblots showed that total ubiquitylation within purified RQC complexes was partially reduced in Rqc1^Ltn1^ co-IPs compared to wild-type control (Fig. 3d). Strikingly, K48-linked polyubiquitylation was virtually undetectable in the mutant (Fig. 3e). The loss of K48-linked chains likely explains why Rqc1 deletions abrogate Cdc48 recruitment to RQC complexes, as previously reported^20,22^. Consistent with this, Cdc48 recovery was greatly reduced in Rqc1^Ltn1^ compared to WT control (Fig. 3f, g).

Similar experiments were performed to test the functional significance of the interaction between Rqc1 and ubiquitin. The predicted structure suggested that E637 and E657 of Rqc1 are key residues for this interaction (Fig. 3c). Based on this, cells were expressed with Rqc1 mutants substituting E637 and E657 with alanines (E637A & E657A, Rqc1^Ub^) in an Rqc1Δ background and used to test whether disrupting this interaction would impair the formation of K48 poly-ubiquitin linkages. Like Rqc^Ltn1^, co-IP of Rqc1^Ub^ showed that the recovery of ubiquitin was diminished compared to wild-type control and that K48-linked chains were not detectable (Fig. 3d, e). As expected, Cdc48 recovery in the Rqc1^Ub^ mutant was also greatly reduced compared to WT control (Fig. 3f, g). The loss of K48-linked polyubiquitin and Cdc48 recruitment to RQC in both Rqc^Ltn1^ and Rqc1^Ub^ mutants prompted us to test their effects on stalled substrate degradation. As expected, the expression of a nonstop mRNA in these cells led to an accumulation of stalled substrate (Fig. 3h). In essence, Rqc1^Ub^ mutations phenocopy Rqc1^Ltn1^, thus indicating that the interactions among Rqc1, Ltn1, and ubiquitin are functionally linked.

The ubiquitin molecule associated with Rqc1 is likely appended to the nascent chain, although we do not observe continuous density of the nascent chain outside of the ribosome. Parts of the Rqc1^CTD^ are positioned within 10 Å of the exit tunnel opening and would overlap with the binding of other ribosome-associated factors that are known to engage nascent chains, such as the nascent polypeptide-associated complex (NAC) and signal recognition particle (SRP)^39^ (Extended Data Fig. 5). The binding of Rqc1 may prevent the association of these factors from interacting with stalled nascent chains.

## Rqc1 facilitates K48-linked polyubiquitylation by Ltn1

Rqc1 facilitates the formation of K48-linked ubiquitin chains by Ltn1 through an unknown mechanism^21^. The interactions among Rqc1^CTD^ with Ltn1^RING^ and ubiquitin rationalize how this occurs. RING domain E3 ligases directly transfer ubiquitin between the ubiquitin-conjugated E2 (E2-Ub, donors) and their substrates (acceptors). The specificity of polyubiquitin linkages requires the proper positioning of the acceptor ubiquitin to the active site of E2-Ub^40,41^. This led us to hypothesize that Rqc1 functions to position the acceptor ubiquitin in a specific orientation that favors K48-linkages with the donor ubiquitin from E2-Ub. We therefore extended our analysis by using AlphaFold3 to generate a predicted structure containing of Rqc1^CTD^, Ltn1^RING^, and the acceptor ubiquitin in complex with E2-Ub (Rad6-ubiquitin). This produced a high-confidence prediction of the complex, and the fitting of the complex into our experimental density containing Rqc1^CTD^, Ltn1^RING^, and acceptor ubiquitin showed that the E2 enzyme and donor ubiquitin were positioned in a solvent-accessible region free of clashes (Fig. 4a and Extended Data Fig. 6). This modeling indicates that the Rqc1-bound acceptor ubiquitin is oriented such that its K48 residue becomes the only accessible lysine proximal to the active site cysteine of the E2 and the C-terminal glycine of the donor ubiquitin (Fig. 4b). The model also agrees with other experimentally determined structures of other RING domains bound to E2-Ub complexes (Extended Data Fig. 6b). Together, these results suggest that Rqc1 holds the acceptor ubiquitin in a defined position to facilitate the specific assembly of K48-linked polyubiquitin chains on stalled nascent chains. As the nascent chain becomes elongated by CAT tails, the ubiquitin chain is presumably pushed toward Cdc48-Ufd1-Npl4, where Npl4 would be in position to unfold the initiator ubiquitin and feed it into the Cdc48 central pore (Fig. 4c). Consistent with this, the distance separating the acceptor ubiquitin on Rqc1 and Npl4 is approximately 50 Å, and it is conceivable that the extension of polyubiquitin chains at ∼25 Å per ubiquitin is sufficient to bridge its gap with Cdc48.

**Fig. 4.**
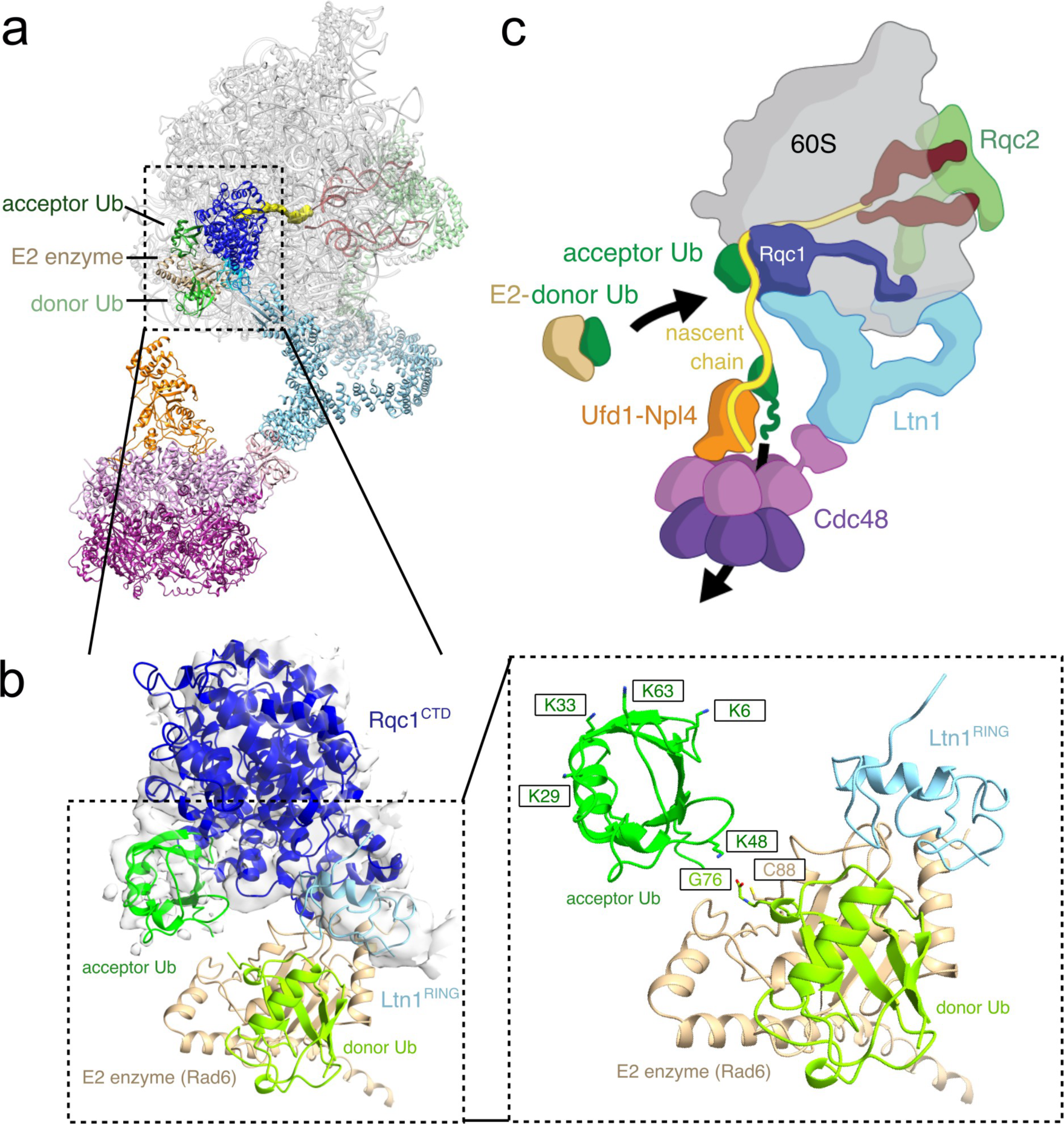
Model of stalled nascent chain extraction by the RQC complex. **a,** Model of the RQC complex including the predicted model of E2 (Rad6) and its accompanying donor ubiquitin. **b,** Close up view of Rqc1^CTD^, acceptor ubiquitin, and Ltn1^RING^ fitted in cryo-EM reconstruction density with the predicted orientation of E2 bound to donor ubiquitin. Key interaction residues between E2, donor ubiquitin (G76), and acceptor ubiquitin (K48) are labeled. The positions of other lysines on the acceptor ubiquitin are also labeled for spatial context. K11 and K27 are also inaccessible for ubiquitylation but are not labeled due to obscured views. **c,** Schematic of nascent chain extension, ubiquitylation, and extraction by the RQC complex.

## Conclusion

Our study provides a comprehensive structure of the RQC complex containing all major components, including Ltn1, Rqc2, Rqc1, and the Cdc48-Ufd1-Npl4 complex. The structure establishes the mechanistic basis for Cdc48 recruitment to RQC substrates via a direct interaction with Ltn1, and this interaction couples substrate ubiquitylation with their extraction from 60S ribosomes. Additionally, Rqc1 plays a critical role in positioning Ltn1^RING^ and ubiquitin to facilitate the formation of K48-linked polyubiquitin chains. The addition of CAT tails by Rqc2 is likely required to extend specific lysine residues to reach the RING domain of Ltn1^13,14^. In yeast, this CAT-tailing process is terminated by the peptidyl-tRNA nuclease Vms1^42–44^. Given that Vms1 interacts directly with Cdc48^45^, it remains to be determined whether this interaction plays a functional role in the RQC pathway. It is also curious why Cdc48-Ufd1-Npl4 is needed to remove the nascent chain, i.e. why the nascent chain does not simply diffuse out of the ribosome following peptidyl-tRNA cleavage. One possibility is that Vms1-mediated cleavage leaves unusual CCA trinucleotide signatures on the nascent chain that may not be readily diffusible through the 60S exit tunnel^46^. An alternate and non-mutually exclusive possibility is that the pulling force from Cdc48 alters the composition of CAT tails to favor nascent chain release^47^.

## Methods

### Cell growth & lysis

Yeast cells (S. cerevisiae strain BY4741) were grown in either rich media or synthetic dropout media and harvested at OD600 of ∼1.2. The cells were pelleted at 4,500 x g for 6 minutes, washed in ice cold water, pelleted again at 4,000 x g for 6 minutes, then resuspended in lysis buffer (50mM HEPES-KOH pH 6.8, 150mM KOAc, 2mM Mg(OAc)_2_, 1mM CaCl_2_, 0.2M sorbitol, and supplemented with cOmplete protease inhibitor cocktail (Roche)). Cell droplets were frozen in liquid nitrogen, lysed into powder using a Freezer/Mill® (SPEX SamplePrep), and stored at -80°C until use. A list of yeast strains used in this study is provided in Extended Data Table 1.

### RQC immunoprecipitation

RQC particles were purified from cells expressing a C-terminal 3xFLAG tag at the endogenous locus of Rqc1, as previously described ^10,22,25^. Lysed yeast powder was thawed on ice and resuspended in immunoprecipitation buffer (95 mM KCl, 5 mM NH_4_Cl, 50 mM HEPES (pH 7.5), 1 mM DTT, 15 mM Mg(OAc)_2_, 0.5 mM CaCl2, supplemented with cOmplete protease inhibitors (Roche)). Supernatant was clarified by centrifugation and incubated with anti-FLAG M2 agarose resin (Sigma) overnight at 4°C. The resin was washed thoroughly with buffer containing ADP-BeFx, and the RQC particles were eluted with excess 3xFLAG peptide for 30 min at 4°C. Purified samples were analyzed by silver stain SDS-PAGE and negative stain electron microscopy and used for cryo-EM experiments and immunoblot analyses.

### Electron microscopy

4 μl of purified RQC particles were cross-linked using 0.15% glutaraldehyde and applied to either Quantifoil R2/2 or R1.2/1.3 grids (SPT Labtech) deposited with a fresh homemade graphene oxide layer, as previously described ^48^. Grids were plunge frozen in liquid ethane using a Mk. II Vitrobot (Thermo Fisher Scientific) with a wait time of 25 s and -1 mm offset. Datasets were collected across six separate sessions on a Titan Krios G3 (Thermo Fisher Scientific) operating at 300kV, equipped with a post-GIF K3 direct detector (Gatan, Inc.). Settings used for data collection are summarized in Extended Data Table 2. Images from five sessions were recorded using SerialEM software^49^ at a nominal magnification of 81,000X in counting mode, corresponding to a pixel size of 1.058 Å with a total dose of approximately 36 electrons/Å², and 40 frames per movie. The sixth session was recorded in super-resolution mode, corresponding to a pixel size of 0.529 Å with a total dose of approximately 50 electrons/Å^2^, and 56 frames per movie. Super-resolution movies were 2x Fourier binned for downstream processing. All movies were set in SerialEM to be recorded between -0.8 to -1.5 μm underfocus with a 3x3 multishot array and either 2 exposures per hole (R1.2/1.3 grids) or 4 exposures per hole (R2/2 grids).

### Image processing and 3D reconstruction

Six datasets were collected and processed. Four datasets were merged into a single group containing 28,302 micrographs, while the remaining two datasets were merged into another group containing 12,667 micrographs. The two groups were processed separately. After data collection, movies were subjected to patch motion correction and patch CTF estimation using cryoSPARC ^50^.

For the first merged dataset, 28,302 micrographs were curated by filtering for exposures with an average defocus between -0.5 and -2.7 μm, CTF fit resolution better than 13.7 Å, and average intensity values ranging from -543 to 346.4. Defocus tilt angles between -9.3 and 28.5 were also included. After manually removing low-quality micrographs, 26,672 micrographs were retained. A total of 1,416,254 particles were initially picked from 5,000 micrographs using a blob picker with a particle diameter constraint to 200–360 Å. Particles were extracted with a box size of 640 pixels and Fourier-cropped to 128 pixels, then subjected to several rounds of 2D classification. High-quality 2D classes were subsequently used as templates for particle picking. After template-based particle picking and manual inspection, 8,338,295 raw particles were extracted with a box size of 640 pixels and Fourier-cropped to 128 pixels for further rounds of 2D classification to exclude non-particles (e.g., non-protein images, ice, and other artifacts). After 2D classification, 699,664 particles were retained and used for ab-initio reconstruction followed by heterogeneous refinement with five classes. One of these classes, consisting of 239,021 particles, contained recognizable RQC features (e.g., Ltn1, Rqc2, and Cdc48) and was subjected to additional 3D classification and heterogeneous refinement. A total of 170,995 particles from this dataset were retained for combination with particles from the second dataset.

For the second merged dataset, micrographs were curated with CTF fit resolution better than 9.7 Å and average intensity values between -399.6 and 191.5, resulting in 12,187 curated micrographs. Template picking was performed using templates derived from the first dataset, resulting in the selection of 1,879,959 particles after manual inspection. These particles were extracted with a box size of 640 pixels and Fourier-cropped to 128 pixels, followed by several rounds of 2D classification, yielding 841,042 particles. These particles underwent heterogeneous refinement with five classes, and one class, containing 332,448 particles with RQC features, was subjected to additional 3D classification and heterogeneous refinement. A total of 260,017 particles were retained and combined with the 170,995 particles from the first dataset. Combined particles underwent two rounds of homogeneous refinement and 3D variability analysis (3DVA)^51^ with a mask surrounding the Ltn1^RING^ and Ltn1^RWD^ domains as well as parts of the 60S ribosomal subunit, with resolution filtered to 15 Å. A 3D variability display was generated using 20 clusters.

A total of 180,785 particles were re-extracted with a box size of 640 pixels and Fourier cropped to 256 pixels. Two additional rounds of homogeneous refinement and 3D variability analysis were performed: the first round applied a mask around the Ltn1^RING^ and Ltn1^RWD^ domains and the 60S subunit, with resolution filtered to 8 Å using 10 clusters; the second round used a mask surrounding the entire structure except for Cdc48, with resolution filtered to 8 Å and 10 clusters. After this step, 159,575 particles were retained for local refinement with a mask around Ltn1 and Rqc1. These particles were re-extracted with a box size of 640 pixels and Fourier-cropped to 320 pixels. Subsequent rounds of non-uniform refinement and 3D classification (10 classes, resolution filtered to 8 Å) yielded 121,159 particles for another round of non-uniform refinement and local refinement, during which particle subtraction was used to remove densities not associated with Ltn1 and Rqc1. After additional 3D classification with 10 classes at 10 Å resolution, particles containing Cdc48 densities were selected. A total of 77,542 particles were extracted with a box size of 640 pixels for non-uniform and local refinement, with a mask around Ltn1 and Rqc1 while subtracting other densities. This produced a density map of Ltn1 and Cdc48 at 6.6 Å resolution.

To focus on the entire complex, the 77,542 particles were recentered on Cdc48 and reextracted with a box size of 256 pixels for 2D classification to obtain high-resolution 2D classes. Selected particles were re-extracted with a box size of 640 pixels and subjected to non-uniform refinement. Finally, 28,262 particles were recentered on the 60S ribosomal subunit, re-extracted with a box size of 640 pixels, and processed through a final round of non-uniform refinement, achieving a 3.2 Å resolution map of the entire complex.

To improve the local resolution of Rqc1^CTD^, a total of 209,041 particles were selected from 3D variability analysis (3DVA) following homogeneous refinement of combined particles. This analysis used a mask encompassing the Ltn1^RING^ and Ltn1^RWD^ domains along with the 60S subunit, with the resolution filtered to 15 Å and divided into 10 clusters. Of these, five clusters exhibited density features consistent with Rqc1 and were subjected to further homogeneous refinement and 3DVA. A mask surrounding Rqc1, the Ltn1^RING^, and the 60S subunit was applied, with resolution filtered to 13 Å across six clusters, retaining 197,360 particles. These particles were re-extracted with a box size of 640 pixels and Fourier-cropped to 320 pixels, then processed through homogeneous refinement and another round of 3DVA using a mask focused on Rqc1, with resolution filtered to 6 Å and analyzed across eight clusters. From these, a single class with 14,039 particles was identified with clear secondary structure density consistent with Rqc1 features. The selected particles were re-extracted with a box size of 640 pixels and subjected to local refinement with a mask around Rqc1, producing a 3.7 Å resolution map.

### Structure prediction

Alphafold3 predictions were carried out through the AlphaFold3 web server (https://golgi.sandbox.google.com/about) ^52^. For the Cdc48 (UniProt P25694) and Ltn1(UniProt Q04781) interaction (fig. S3A), the input sequence for Cdc48 was L36-G196, and the input sequence for Ltn1 was L933-E1074. For the Rpl38 (UniProt P49167) and Rqc1 (UniProt Q05468) interaction, the input sequence for Rpl38 was M1-L78, and the input sequence for Rqc1 was D120-Y143 (fig. S4A).

For the interaction between the Ltn1 RING domain (UniProt Q04781) and Rqc1 (UniProt Q05468), the input sequence for Ltn1’s RING domain was E1507-R1562, and the input sequence for Rqc1 was G144-G723 (fig. S4a). For the interaction between ubiquitin (UniProt P0CG63) and Rqc1 (UniProt Q05468), the input sequence for ubiquitin was M1-G76, and the input sequence for Rqc1 was G144-G723 (fig. S4A).

For the interactions among Ltn1 RING domain (UniProt Q04781), donor and acceptor ubiquitins (UniProt P0CG63), Rad6 (UniProt P06104), and Rqc1(UniProt Q05468), the input sequence for Ltn1’s RING domain was E1507-R1562, for Ubiquitin was M1-G76, for Rad6 was M1-D172, and for Rqc1 was G144-G723 (fig. S6A).

### Model fitting and structure visualization

Rigid body fittings were done in UCSF Chimera ^53^ or UCSF ChimeraX ^54^. Extraribosomal densities were distinguished by computing difference maps between cryoEM reconstructions (EMDB entries EMD-15431, EMD-15433 & EMD-15434) and reference yeast 60S maps from crystal structure coordinates (PDB entry 8AGW) ^11^.

For the Cdc48-Ltn1 interface, the predicted structural model consisting of the N-terminal domain of Cdc48 and the elbow region of Ltn1 was fitted as a rigid body in ChimeraX. For fitting of the Cdc48 hexamer, the structure of the Cdc48-Ufd1/Npl4 complex with the Cdc48NTD in the ‘up’ conformation was also fitted as a rigid body in ChimeraX (PDB ID: 8DAU ^55^). For the subcomplex containing Rqc1CTD, acceptor Ub, E2 (Rad6), donor Ub, and Ltn1RING, we manually guided the fitting of Ltn1RING into its density, and then used UCSF Chimera to refine the rigid body fitting.

All figures were prepared with UCSF Chimera ^53^, UCSF ChimeraX ^54^, Pymol (https://www.pymol.org), and intermolecular interactions were analyzed using PDBePISA ^56^.

### Protein A-Nonstop substrate degradation/accumulation assay

The Protein-A nonstop plasmid (pAV184) was generously provided by Ambro van Hoof^34^. Yeast was grown to saturation overnight in the corresponding selection SRaf media, then back diluted to an OD600 of approximately ∼0.15 and grown for 3-4 hours to reach early log phase (A600 ∼0.3–0.5). At this point, 1% galactose was added to induce expression of the GAL1 promoter for 3 hours and then harvested. Cells were lysed by boiling for 5∼10 minutes in Laemmli buffer.

### Immunoblots

Samples were separated by SDS-PAGE, followed by electrophoretic transfer onto polyvinylidene difluoride membrane (Bio-Rad). Membranes were blocked with 5% nonfat milk in TBST (20 mM Tris-HCl, pH 8.0, 150 mM NaCl, 0.1% Tween-20) for 30-60 min at room temperature.

For general ubiquitin and anti-K48 linkage-specific ubiquitin immunoblots, yeast cells expressing a stalling substrate reporter were used. The reporter contains a 12x polyarginine, including pairs of the difficult-to-decode CGA codon, inserted between the coding regions of GFP and RFP, as previously used (*10*). The membrane was incubated overnight at 4o C with the anti-K48-linkage specific ubiquitin or anti-ubiquitin at 1:5,000 dilution. Membranes were washed in TBST, followed by incubation with secondary antibody, donkey *α*-rabbit conjugated to horseradish peroxidase (HRP, GE Biosciences; 1:10,000 dilution). Antibodies were detected using the SuperSignal West Dura Chemiluminescent Substrate (Thermos Scientific). The membranes were thoroughly washed with TBST and incubated for 1h at room temperature with anti-FLAG clone M2 (ABclonal; 1:5,000 dilution). Membranes were washed in TBST, followed by incubation with secondary antibody IRDye 680RD Goat Anti-Rabbit (LI-COR 926-68071; 1:20,000 dilution) for 1 h at room temperature.

For anti-Cdc48 and anti-FLAG immunoblots, the membranes were incubated for 1h at room temperature with anti-Cdc48 (gift from Thomas Sommer; 1:5,000 dilution). Membranes were washed in TBST, followed by incubation with secondary antibody IRDye 680RD Goat AntiRabbit (LI-COR 926-68071; 1:20,000 dilution) for 1 h at room temperature. The membranes were thoroughly washed with TBST and incubated for 1h at room temperature with anti-Flag (Sigma; 1:5,000 dilution). Membranes were washed in TBST, followed by incubation with secondary IRDye 800CW goat anti-mouse (LI-COR 926-32210; 1:10,000 dilution) for 1 h at room temperature. Membranes were washed in TBST and digitized using an Odyssey CLx scanner (LI-COR).

For immunoblots against nonstop Protein A substrate, cells were expressed with the pAV184 plasmid as described above. The membrane was incubated for overnight at 4o C or 1 hour at room temperature with anti-protein A antibody (Sigma; 1:10,000 dilution). Membranes were washed in TBST, followed by incubation with secondary antibody, donkey *α*-rabbit conjugated to horseradish peroxidase (HRP, GE Biosciences; 1:10,000 dilution). Antibodies were detected using the SuperSignal West Dura Chemiluminescent Substrate (Thermos Scientific). The membranes were thoroughly wash with TBST and incubated for 1h at room temperature with anti-Pgk1 (Thermos Scientific; 1:5,000 dilution). Membranes were washed in TBST, followed by incubation with secondary IRDye 800CW Goat Anti-Mouse (LI-COR 926-32210; 1:10,000 dilution) for 1 h at room temperature. Membranes were washed in TBST and digitized using an Odyssey CLx scanner (LI-COR).

## Data availability

The cryo-EM maps of RQC complex are deposited to the Electron Microscopy Data Bank with the accession codes EMD-48531 (consensus reconstruction with Cdc48 density), EMD-48529 (Ltn1-Cdc48 focused reconstruction), and EMD-48530 (Rqc1 focused reconstruction).

## Acknowledgments

We thank Ambro van Hoof for providing the Protein A-nonstop reporter plasmid (pAV184) and Thomas Sommer for the anti-Cdc48 antibody. We thank David Belnap and the University of Utah Arnold and Mabel Beckman Center for CryoEM for cryo-EM support and the University of Utah Center for High Performance Computing for computational support.

## Author contributions

Methodology: WL, TC, PSS. Investigation: WL, TC, PSS. Visualization: WL, PSS. Supervision: PSS. Writing – original draft: WL, PSS. Writing – review & editing: WL, PSS. Funding acquisition: PSS. Project administration: PSS.

## Competing interests

The authors declare that they have no competing interests.

## Funding

National Institutes of Health grant R35 GM133772 (PSS)

## Notes

### Competing Interest Statement

The authors have declared no competing interest.

