## supplemental file for "Mechanism of nascent chain removal by the Ribosome-associated Quality Control complex"

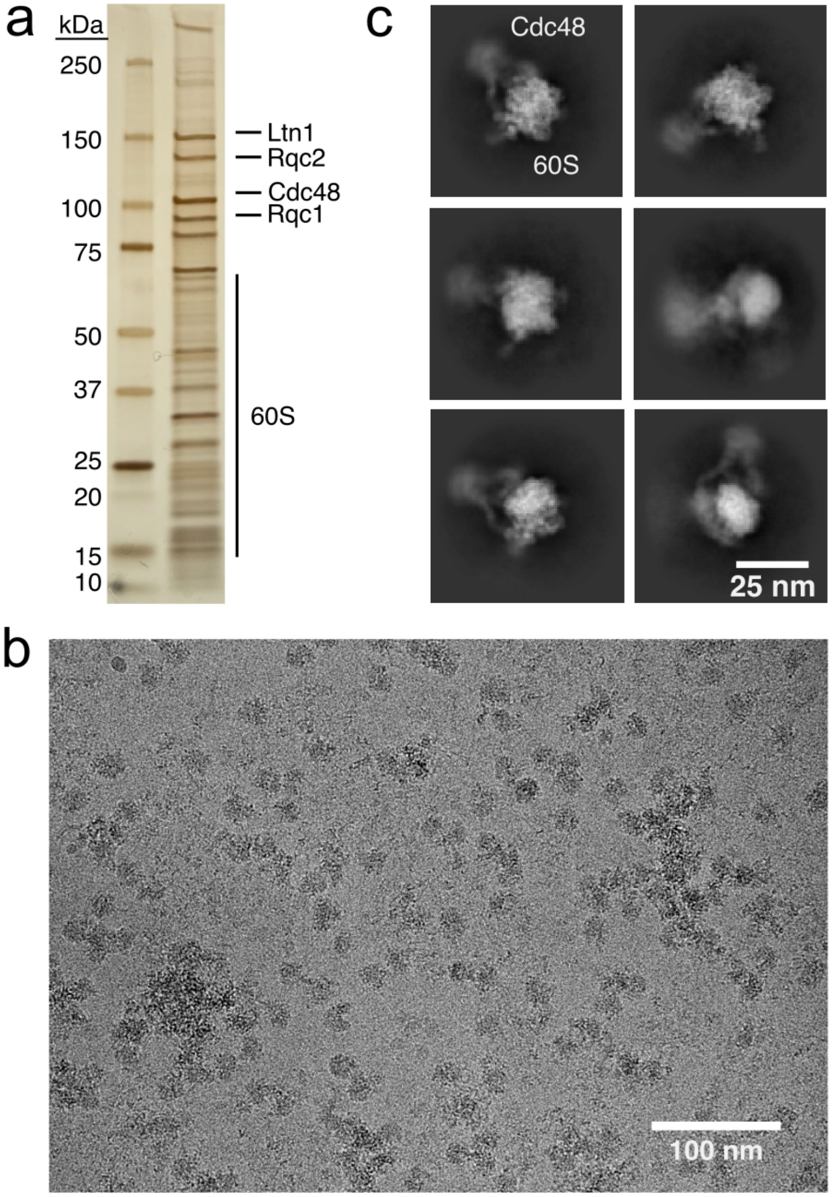


**Extended Data Fig. 1. Co-immunoprecipitation and Cryo-EM structural analysis of the RQC complex. (a)** Silver stained RQC sample following αFLAG (Rqc1-3xFLAG) co-immunoprecipitation. Relevant protein bands labeled. **(b)** Cryo-EM micrograph of purified RQC particles. (**c**) Representative 2D classes of RQC particles.

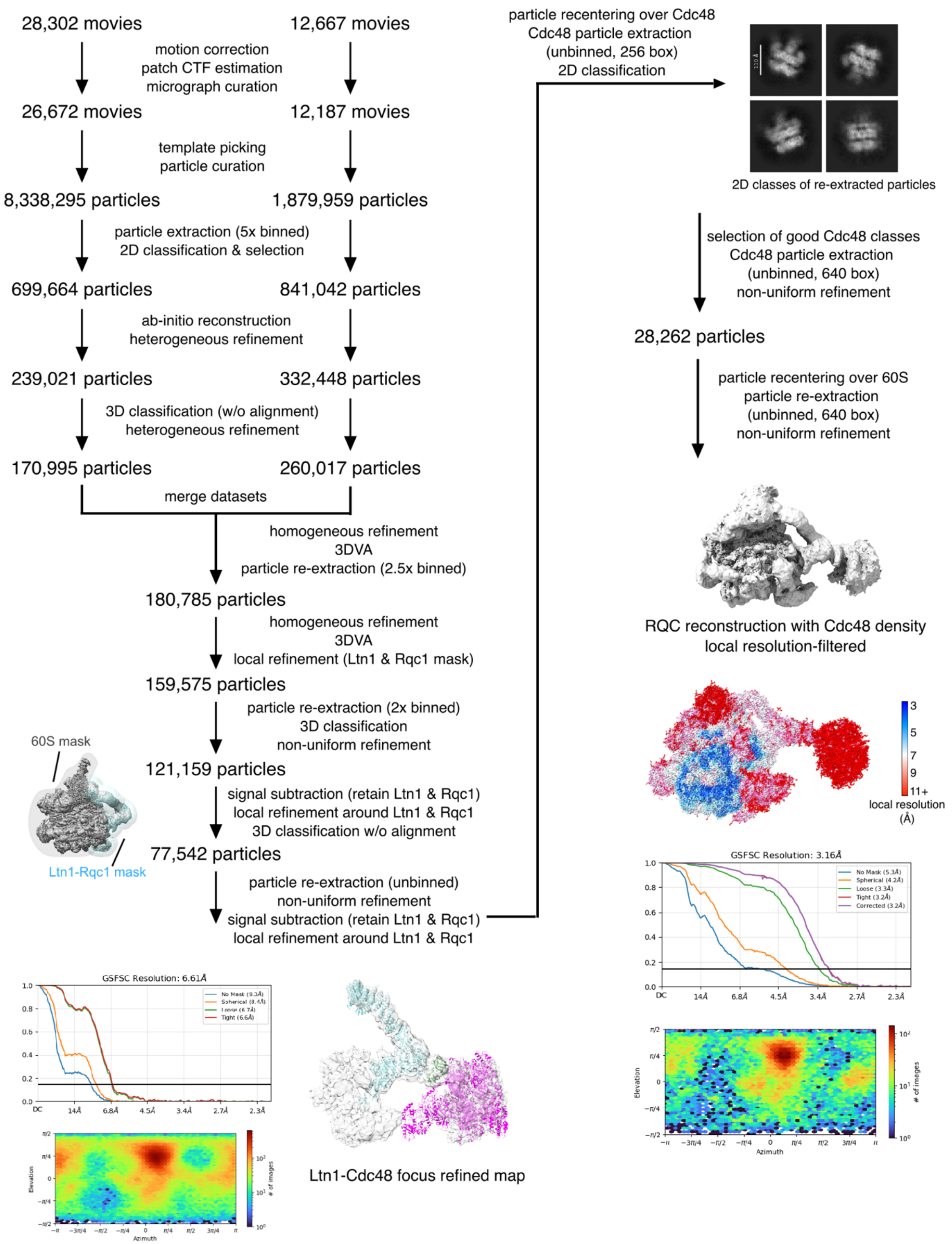


**Extended Data Fig. 2. Cryo-EM Data Processing Workflow.** Schematic of cryo-EM data processing from movies to reconstructions.

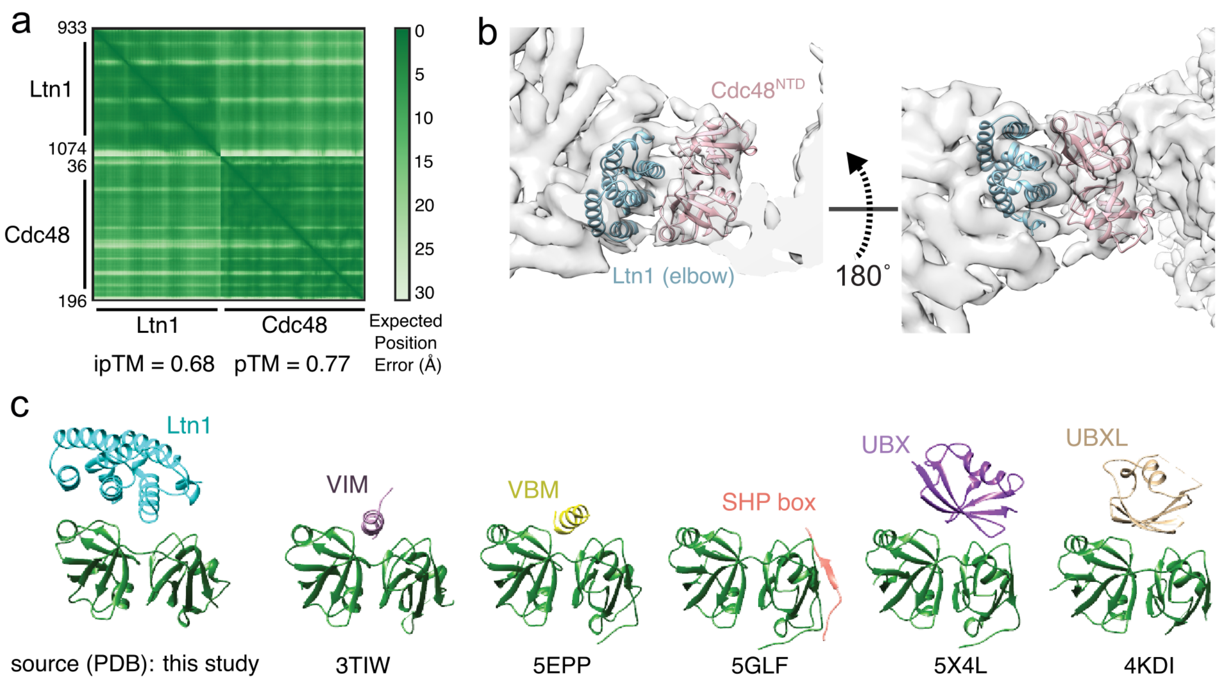


**Extended Data Fig. 3. Cdc48$^{NTD}$ interacts with the Ltn1 elbow. (a)** AlphaFold3 matrix showing the expected position error for each residue in the sequence for the Ltn1 elbow region (residues 933- 1074) and Cdc48NTD (residues 36-196). **(b)** Rigid-body fitting of the predicted Ltn1-Cdc48 model from panel A into the locally refined reconstruction density. **(c)** Ribbon representation of the different types of cofactors binding to the Cdc48/p97 NTD (green).

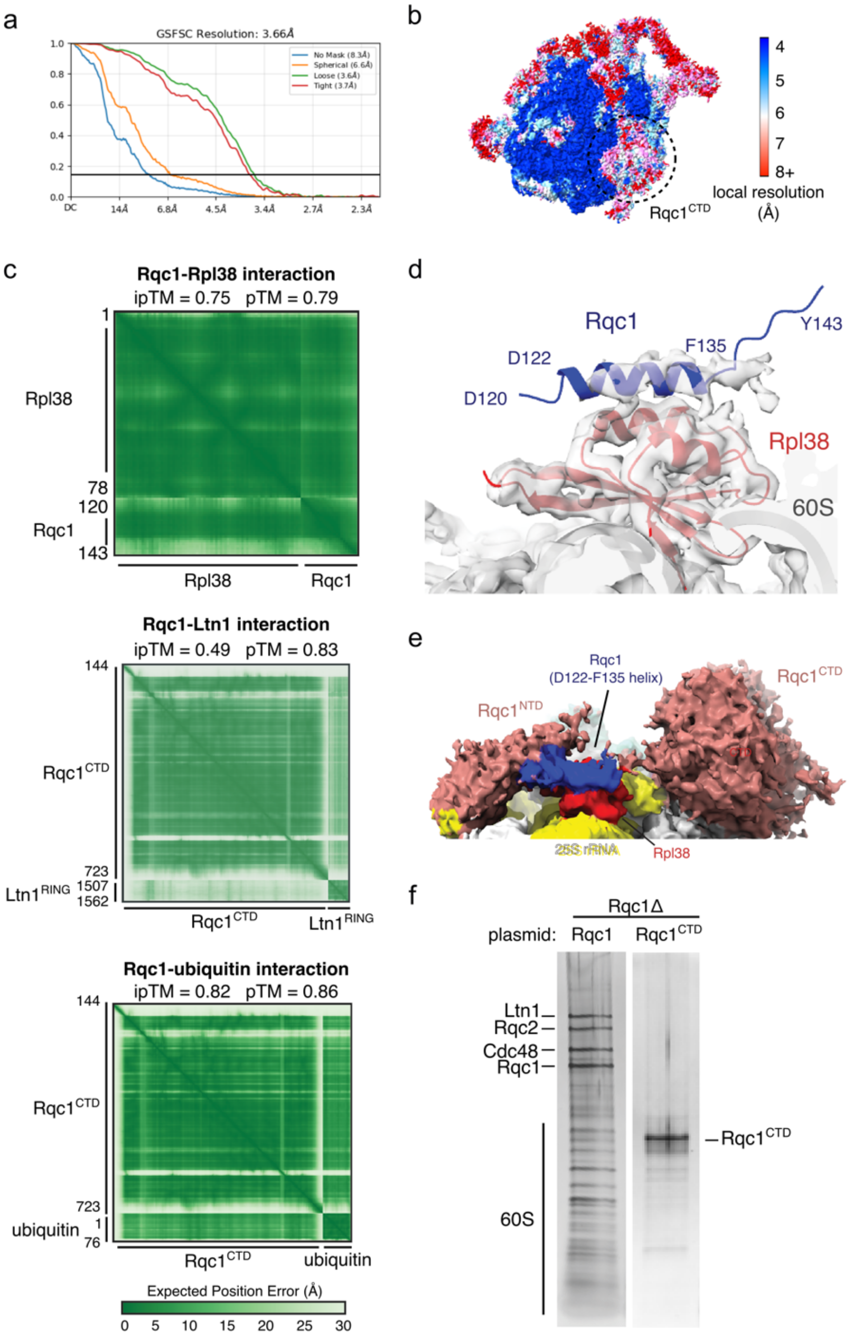


**Extended Data Fig. 4. Rqc1 interacts with the 60S ribosome and Ltn1. (a)** Fourier shell correlation (FSC) plot for the local refinement of Rqc1. **(b)** Local resolution heat map of the Rqc1 local refinement. **(c)** The AlphaFold matrix showing the expected position error for each residue in the predicted structures of Rqc1-Rpl38 (top), Rqc1-Ltn1 (middle), and Rqc1-ubiquitin (bottom). **(d)** The predicted structure of Rqc1-Rpl38 was fitted as a rigid body into the reconstruction density of Rpl38. **(e)** The Rqc1 internal helix (D122-F135) separates the NTD from the CTD. **(f)** Expression and co-IP of Rqc1CTD in Rqc1Δ cells fails to recover ribosomes and other RQC factors.

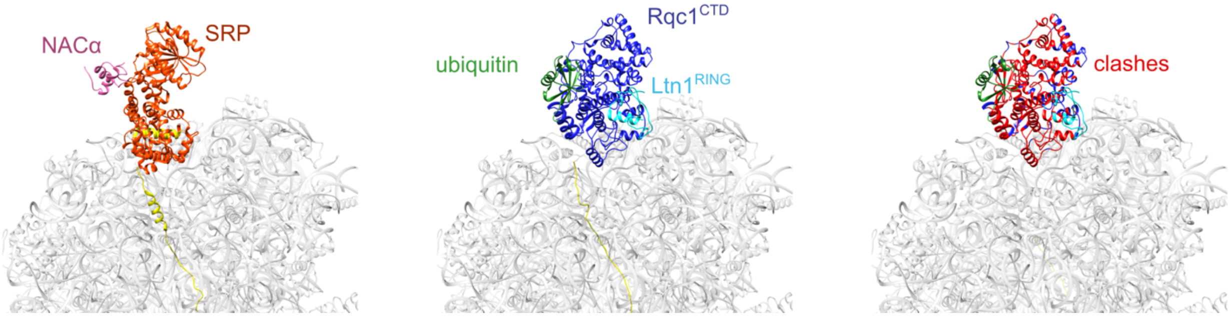


**Extended Data Fig. 5. Rqc1 occupies the same binding site as NAC and SRP.** Left, model of SRP and NACα bound to nascent chain emerging out of the ribosome exit tunnel. Middle, model of Rqc1$^{CTD}$-Ltn1$^{RING}$-ubiquitin bound at the ribosome exit tunnel opening. Right, clashes between Rqc1$^{CTD}$-Ltn1$^{RING}$-ubiquitin and SRP-NACα are indicated in red ribbon.

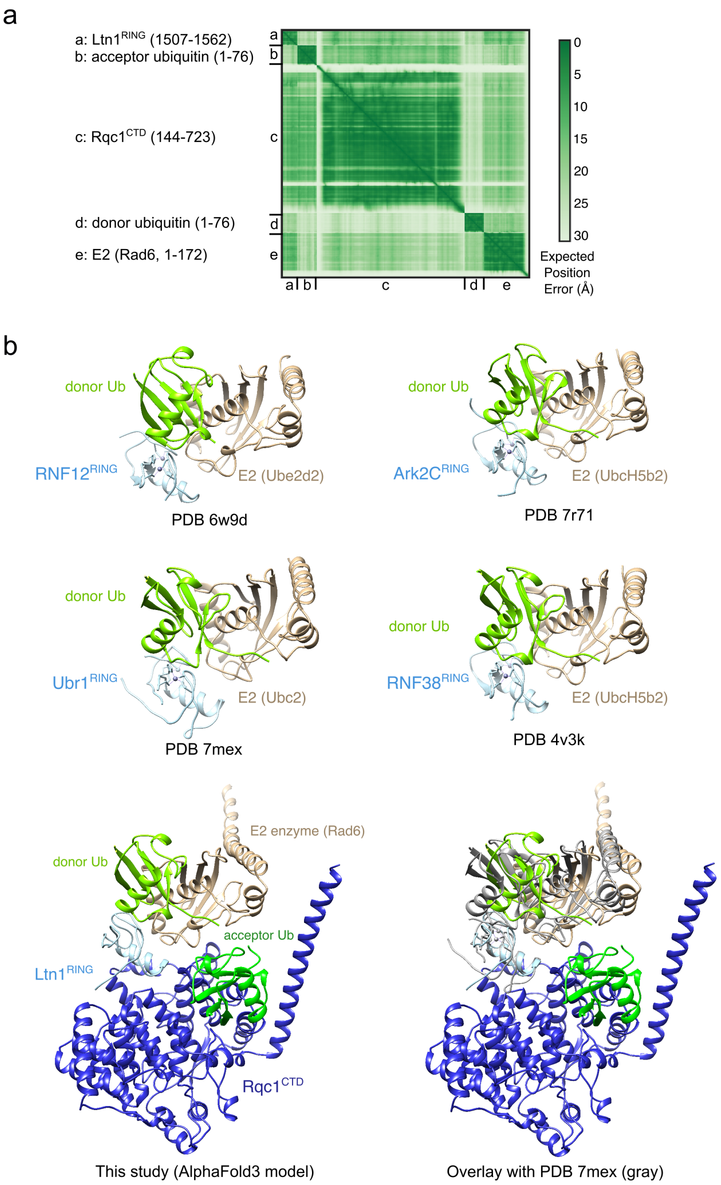


**Extended Data Fig. 6. Rqc1 facilitates K48-polyubiquitylation by Ltn1. (a)** The AlphaFold matrix showing the expected position error for each residue in the sequence for Rqc1CTD, Ltn1RING, acceptor and donor ubiquitins, and Rad6 E2 enzyme. **(b)** Models of E2-donor ubiquitins bound to RING domains of RNF12, Ark2C, Ubr1, and RNF38 are structurally similar to E2-Ub bound to Ltn1. The positioning of Rqc1CTD holds the acceptor ubiquitin in a defined orientation to facilitate K48-linkage.

### Extended Data Table 1. Yeast strains used in this study

| Strain | Background | Plasmid 1 | Plasmid 2 |
|---|---|---|---|
| yPS1253 | BY4741 *rqc1-3xFLAG::kan* | | |
| yWY1001 | BY4741 *rqc1::nat* | pRQC1_Rqc1$^{CTD}$, LEU2 marker | |
| yWY1002 | BY4741 *rqc1::nat ltn1::hyg* | pRQC1_Rqc1$^{CTD}$, LEU2 marker | |
| yWY1003 | BY4741 *rqc1::nat* | pRQC1_WT, LEU2 marker | GFP_TEV_R4_RFP, URA3 marker |
| yWY1003 | BY4741 *rqc1::nat* | pRQC1_ Rqc1$^{Ltn1}$, LEU2 marker | GFP_TEV_R4_RFP, URA3 marker |
| yWY1004 | BY4741 *rqc1::nat* | pRQC1_Rqc1$^{Ub}$, LEU2 marker | GFP_TEV_R4_RFP, URA3 marker |
| yWY1005 | BY4741 *rqc1-3xFLAG::kan ltn1::his met2::pNOP1-hCas9:nat* | pLTN1_Ltn1_WT, LEU2 marker | GFP_TEV_R4_RFP, URA3 marker |
| yWY1006 | BY4741 *rqc1-3xFLAG::kan ltn1::his met2::pNOP1-hCas9:nat* | pLTN1_Ltn1$^{AAA}$, LEU2 marker | GFP_TEV_R4_RFP, URA3 marker |
| yWY1007 | BY4741 *rqc1::nat* | pAV184, URA marker | |
| yWY1008 | BY4741 *rqc1-3xFLAG::kan ltn1::his met2::pNOP1-hCas9:nat* | pAV184, URA marker | |
| yWY1009 | BY4741 *rqc1::nat* | pAV184, URA marker | pRQC1_WT, LEU2 marker |
| yWY1010 | BY4741 *rqc::nat* | pAV184, URA marker | pRQC1_Rqc1$^{Ltn1}$, LEU2 marker |
| yWY1011 | BY4741 *rqc1::nat* | pAV184, URA marker | pRQC1_Rqc1$^{Ub}$, LEU2 marker |
| yWY1012 | BY4741 *rqc1-3xFLAG::kan ltn1::his met2::pNOP1-hCas9:nat* | pAV184, URA marker | pLTN1_Ltn1_WT, LEU2 marker |
| yWY1013 | BY4741 *rqc1-3xFLAG::kan ltn1::his met2::pNOP1-hCas9:nat* | pAV184, URA marker | pLTN1_Ltn1$^{AAA}$, LEU2 marker |

**Extended Data Table 2. Data collection and refinement statistics**

| | RQC consensus map (EMD-xxxx) | Ltn1-Cdc48 focused map (EMD-xxxx) | Rqc1$^{CTD}$-focused map (EMD-xxxx) |
|---|---|---|---|
| **Data collection** | | | |
| Microscope | Titan Krios G3 | Titan Krios G3 | Titan Krios G3 |
| Magnification | 81,000x | 81,000x | 81,000x |
| Voltage (kV) | 300 | 300 | 300 |
| Electron exposure ($e^-$/Å$^2$) | 16.1 | 16.1 | 16.1 |
| Defocus range (μm) | -0.8 to -1.5 | -0.8 to -1.5 | -0.8 to -1.5 |
| Pixel size (Å) | 0.529 | 0.529 | 0.529 |
| Detector | Gatan K3 | Gatan K3 | Gatan K3 |
| Data collection software | SerialEM | SerialEM | SerialEM |
| Total number of frames | 40/56 | 40/56 | 40/56 |
| **Data processing** | | | |
| Number of micrographs | 40,969 | 40,969 | 40,969 |
| Symmetry imposed | C1 | C1 | C1 |
| Final particle images | 28,262 | 77,542 | 14,039 |
| Map resolution (Å) | 3.2 | 6.6 | 3.7 |
| FSC threshold | 0.143 | 0.143 | 0.143 |
